# Rapid Mechanistic Bridging of an Alzheimer’s Disease Plasma Protein Staging Panel Across Brain Proteomics Cohorts

**DOI:** 10.64898/2026.08.31.748393

**Authors:** Charles C. Kim, Meghan Kerrisk Campbell, Maximilien Burq, Aljaz Potocnik, Dejan Stepec, Cássia Maués Cuóco, Yelena Bronevetsky, Domen Zafred, Jacqueline M. Leung, Peter Cimermancic

## Abstract

**Background:** Blood-based biomarkers are transforming Alzheimer’s Disease (AD) diagnosis and staging. Recent multi-protein plasma panels accurately identify individuals with advanced Braak pathology, but whether these circulating biomarkers truly reflect the molecular remodeling underlying AD neuropathology remains unclear. Pre-existing datasets could answer such questions, but their reuse requires harmonized protein expression data, standardized sample annotations, and consistent study metadata.

**Methods:** A recently reported seven-protein plasma staging panel was evaluated across multiple post-mortem proteomics brain cohorts using pre-structured datasets on the Tesorai platform. Five datasets with Braak stage data were identified and re-analyzed. A linear model was used to distinguish late (V-VI) from early Braak stage (0-IV). Because phosphorylated tau 217 (p-tau217) and amyloid beta 1-40 (Aβ40) were not reported for any studies, raw spectra were reprocessed to quantify these proteoforms.

**Results:** Across the cohorts, the plasma-derived biomarker panel consistently discriminated early from late disease despite heterogeneous protein coverage. Reprocessing of raw spectra recovered tau phosphopeptides absent from the original protein summaries, enabling inclusion of p-tau217, while Aβ40 remained undetected. Together, these findings show that proteins comprising a recently proposed blood-based staging panel are associated with proteomic remodeling in the AD brain across independent cohorts.

**Conclusions:** These findings provide biological validation for a recently proposed blood-based seven-protein staging panel by demonstrating that its constituent biomarkers are associated with disease-stage proteomic changes in AD brain tissue. More broadly, re-mining legacy mass spectrometry data for disease-relevant proteoforms, combined with pre-structured datasets, can accelerate evaluation of emerging blood-based biomarkers against neuropathological and molecular features of disease.

## Introduction

Alzheimer’s disease (AD) is defined neuropathologically by the accumulation and regional progression of β-amyloid (Aβ) plaques and hyperphosphorylated tau neurofibrillary tangles, accompanied by progressive synaptic dysfunction, neuronal injury, and ultimately cognitive decline. The topographic distribution of neurofibrillary pathology forms the basis of the Braak staging system, which distinguishes early involvement of transentorhinal and limbic regions (Braak 0–IV) from advanced neocortical involvement (Braak V–VI).^1,2^ Increasing evidence indicates that the extent and anatomical distribution of tau pathology are closely related to clinical progression, with higher tau burden associated with more extensive cognitive decline.^3^ Thus, determining the stage of tau pathology is increasingly important not only for understanding disease biology, but also for prognosis, clinical trial patient selection, and disease activity monitoring.

Historically, characterization of AD pathology in living patients has relied on cerebrospinal fluid (CSF) biomarkers and positron emission tomography (PET) imaging. These approaches provide valuable information on Aβ and tau pathology, but have been limited in their broader clinical and research use due to their cost, availability, procedural complexity, and invasiveness. The rapid development of blood-based biomarkers (BBMs), particularly plasma phosphorylated tau (p-tau), offers a more accessible and scalable approach.^4,5^ Plasma p-tau217 is among the most extensively validated BBMs, demonstrating strong associations with AD pathology and high accuracy for distinguishing biologically defined AD from other neurodegenerative conditions.^6–15^ These findings, in concert with others, contributed to a major milestone in 2025, when the U.S. Food and Drug Administration (FDA) cleared Lumipulse G as the first blood-based diagnostic test for AD, based on the ratio of p-tau217/Aβ1-42.^16^ Even more recently, a blood test based on p-tau217 alone was cleared by the FDA for determining the presence or absence of amyloid pathology.^17^ Despite this progress, p-tau217 is currently used as a readout for early amyloid-associated disease biology rather than advanced neocortical tau burden, underscoring the need for novel BBMs that can accurately stage tau pathology to support clinical trials and patient management.

A recent multicenter study directly addressed these limitations by combining p-tau217 with additional plasma proteins to identify advanced tau pathology.^18^ A seven-protein panel was identified that distinguished individuals with tau PET involvement corresponding to Braak V–VI from those with earlier Braak 0–IV pathology. The multiprotein model achieved AUCs of approximately 0.92–0.94 across discovery and validation cohorts, which was a significant improvement over p-tau217 alone. Together, these findings suggest that circulating proteins beyond p-tau217 may encode information about the biological transition toward widespread neocortical tau pathology.

Although some of the proteins in the plasma panel have previously been implicated in AD pathogenesis, the associations of others are understudied. We sought to address the molecular link between changes in the plasma proteome to remodeling directly within the brain.

Leveraging a database of pre-structured public datasets available on the Tesorai platform, we reanalyzed five post-mortem brain proteomics cohorts with accompanying Braak-stage annotations, including reprocessing of the raw data to identify p-tau217. The results provide mechanistic validation of the recently proposed blood-based staging signature at the molecular level in AD brain tissue and highlight how previously generated proteomics datasets can be efficiently and rapidly re-mined to recover disease-relevant proteoforms.

## Methods

### Dataset preparation for validation of the AD multi-protein panel

Public proteomics datasets remain highly heterogeneous in file structure, protein identifiers, abundance representations, and metadata, limiting their reuse for large-scale computational analyses and requiring substantial study-specific curation and harmonization. To address these challenges, we developed a semi-automated pipeline combining AI-assisted dataset parsing with expert review and, where necessary, reanalysis of raw mass spectrometry (MS) data using Tesorai Search.^19^ Using this workflow, we assembled a harmonized corpus of 440 public MS proteomics datasets comprising 48,843 samples across diverse tissues, diseases, and perturbation studies (**Figure 1**). These structured datasets are available on the Tesorai platform through the Trove database.

**Figure 1.**
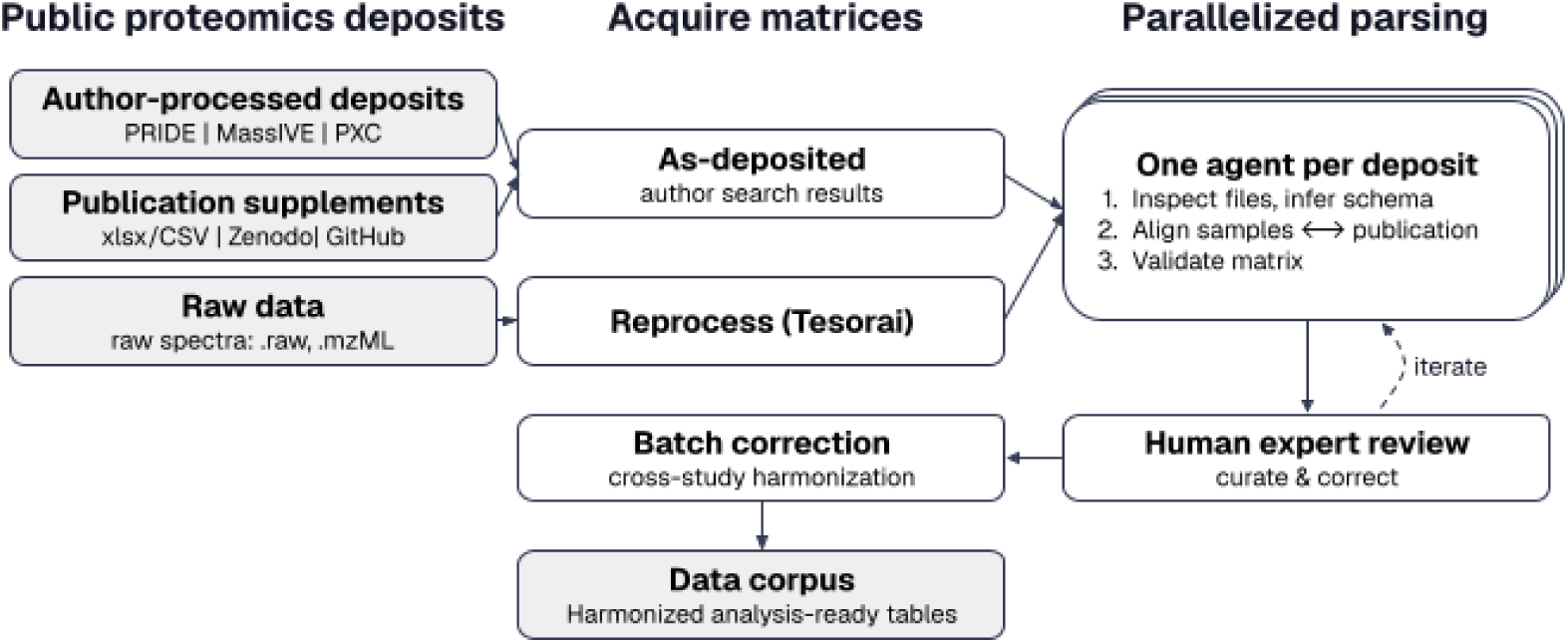
Tesorai’s dataset structuring pipeline. Diverse public deposits (repository search results, publication supplements, and raw spectra) are either taken as-deposited or reprocessed through the Tesorai platform to generate quantified protein matrices. One LLM agent per deposit infers each layout and aligns samples to the source publication; human experts review and iterate with the agents; the harmonized, batch-corrected tables are made available for analysis through a chat interface.

Datasets were selected from the Tesorai Trove database using predefined inclusion criteria. Eligible studies included >20 participants, reported neuropathological Braak staging, employed exploratory (discovery) MS proteomics, and detected at least two of the seven panel proteins (NPTXR, SFRP1, APP as an Aβ40 surrogate, IL-13, GDNF, KLK6, and MAPT as a p-tau217 surrogate). Studies lacking Braak stage annotations or consisting exclusively of targeted proteomic measurements were excluded. Applying these criteria, we identified six independent studies for downstream analyses. To ensure data integrity, we attempted to reproduce the differential expression analyses reported in each of the six source publications and excluded datasets for which the published results could not be reproduced. This quality-control step excluded one dataset, resulting in five studies being retained for downstream analyses (**Table 1**, **Figure 2**).

**Figure 2:**
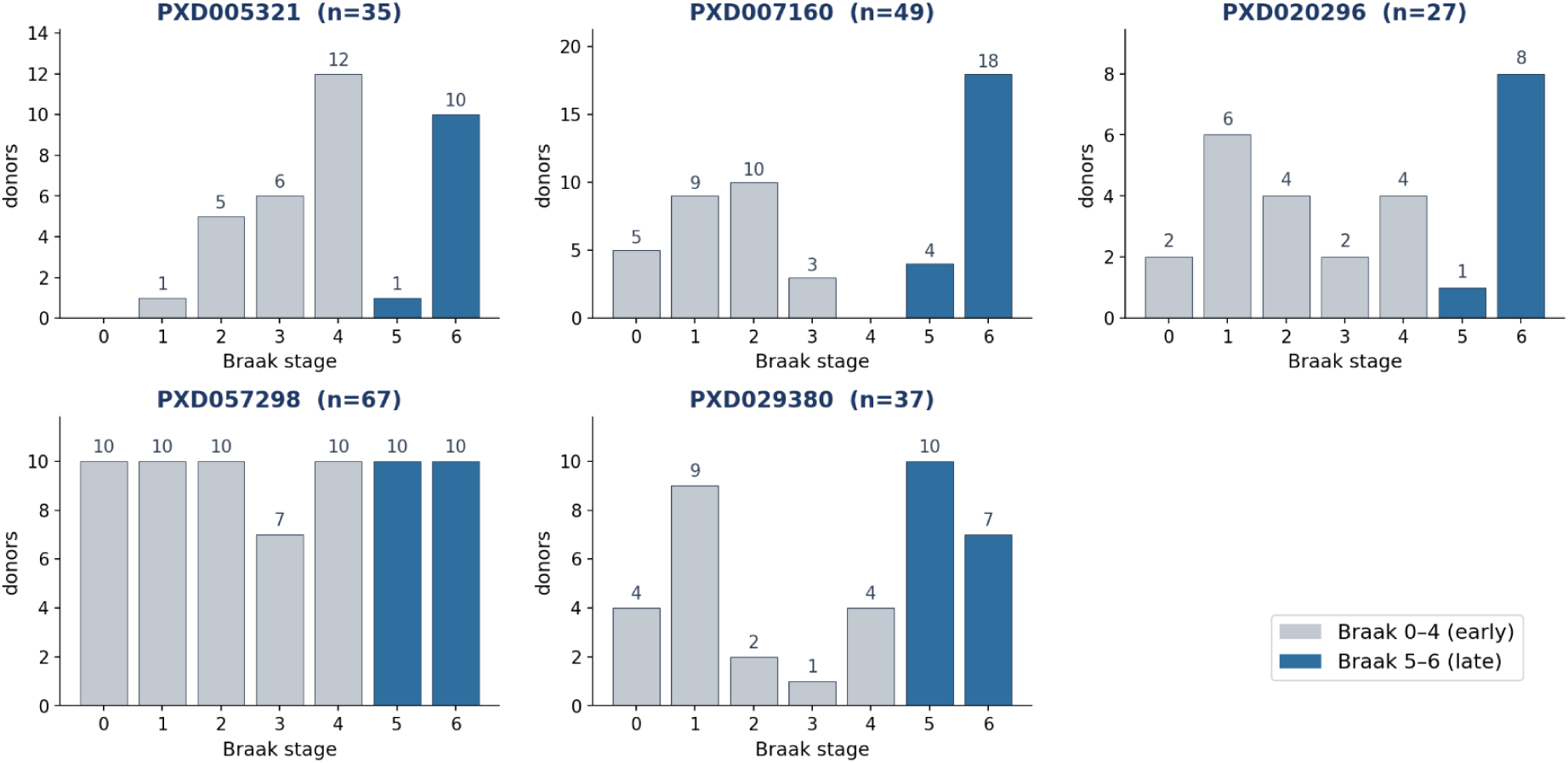
Braak-stage distribution of donors in each proteomics cohort. Bars show the number of donors at each neurofibrillary-tangle (NFT) Braak stage (0–6) per study, grouped into early-stage (Braak 0–4, grey) and late-stage (Braak 5–6, blue) as used in the classification analyses; Braak 0 corresponds to cognitively normal controls. Due to the absence of reported Braak stages for 5 specimens in PXD057298, the total count here is lower than the figure listed in Table 1.

**Table 1.** Public AD proteomics datasets selected for validation of the plasma multi-protein staging panel. Studies were selected from the Trove database based on the availability of AD samples, neuropathological Braak staging, sufficient cohort size, and discovery (untargeted) MS proteomics. The table summarizes the tissue analyzed, total number of samples (S#), number of study participants (P#), and the subset of proteins from the seven-protein plasma staging panel detected in each dataset. Variable panel coverage across studies reflects differences in experimental design and acquisition methodology.

| PXD ID | Publication | S# | P# | Proteins detected from seven-protein panel |
| --- | --- | --- | --- | --- |
| PXD005321 | Detergent-insoluble brain proteome linked to amyloid and tau in AD.<sup>20</sup> | 35 | 35 | 2/7: MAPT, APP |
| PXD007160 | Global quantitative analysis of the human brain proteome in AD and PD.<sup>21</sup> | 80 | 49 | 5/7: MAPT, NPTXR, SFRP1, APP, KLK6 |
| PXD020296 | Global brain proteome and phosphoproteome in AD.<sup>22</sup> | 27 | 27 | 4/7: MAPT, NPTXR, APP, KLK6 |
| PXD057298 | Extracellular exosome signature links hypertension and AD.<sup>23</sup> | 72 | 72 | 5/7: MAPT, NPTXR, SFRP1, APP, KLK6 |
| PXD029380 | Proteomic profiling in cerebral amyloid angiopathy (HTRA1).<sup>24</sup> | 37 | 37 | 4/7: MAPT, NPTXR, SFRP1, APP |

### Study-specific normalization and batch correction

#### PXD007160 (TMT10, SPS-MS3, offline high-pH reversed-phase fractionation)

Reporter-ion intensities were extracted at the PSM level and normalized to the pooled global internal standard (GIS) channels (126 and 131) within the corresponding TMT plex. This within-plex calibration was performed before aggregating peptide measurements to proteins across fractions. GIS-normalized protein abundances were log2-transformed to harmonize measurements across the ten TMT plexes. For descriptive analyses, we additionally applied ComBat using plex as the batch variable.^25^ Fraction was not modeled as a batch effect because fractions were nested within plex and shared across reporter channels rather than representing a between-sample source of variation. ComBat reduced the variance attributable to plex structure in PC1 from 12.9% to 0.04%. Because GIS normalization is performed independently of other biological samples, it was used directly for cross-validated analyses (**Figure S1**). In contrast, because ComBat estimates batch parameters across samples, ComBat-corrected data were used only for descriptive analyses unless batch correction was fitted exclusively within the training folds.

#### PXD020296 (TMT11, MS2, high-pH fractionation; three plexes)

We used the authors’ deposited GIS-normalized and centered peptide matrix, with peptide measurements subsequently aggregated to the protein level across fractions. Thus, per-plex reference-channel calibration and cross-fraction aggregation had already been performed. As in PXD007160, plex represented the relevant multiplexing unit for batch correction. Residual plex-associated structure was limited, accounting for 6.7% of PC1 variance, and decreased to 0.9% following optional ComBat correction using plex as the batch variable. This correction did not materially affect downstream results. This dataset additionally included IMAC phospho-enriched acquisitions (36 enriched versus 72 global raw files across the three plexes); protein-level quantification was derived exclusively from the global total-proteome runs, with the phospho-enriched runs excluded.

#### PXD057298 (label-free DIA, timsTOF, single-shot DIA-PASEF)

We used the authors’ DIA-NN protein quantification matrix and applied a log2 transformation before downstream analysis. This experiment consisted of single-injection DIA-PASEF measurements without peptide prefractionation or multiplexed reporter channels. As no sample preparation batch variable was documented, no additional batch correction was applied.

#### PXD029380 (label-free DDA, MaxQuant/MaxLFQ, single-shot per sample)

We used the authors’ MaxQuant LFQ intensities, which were jointly normalized across runs during MaxLFQ processing, and applied a log2 transformation for downstream analysis. Samples were analyzed as single injections without prefractionation. No batch variable was documented, so no batch correction was applied.

#### PXD005321 (label-free XIC, DQuan; gel-based fractionation into run sets)

We used the deposited protein-level log2 XIC abundance matrix, which had already undergone DQuan normalization, including cross-gel normalization using shared controls. No additional transformation was applied. Samples were distributed across two gel run sets (Gel 1 and Gel 2), representing the only identifiable potential batch variable. However, gel assignment was unavailable for all donors, including control samples, so we did not apply ComBat or other explicit batch correction and analyzed the deposited normalized matrix as provided.

Because each study was quantified and normalized using its platform-specific measurement scale, including TMT reporter-ion ratios, MaxLFQ intensities, DIA-NN protein quantities, and XIC-based abundances, absolute values were interpreted within, rather than across, studies. Cross-study comparisons therefore focused on predictive performance, standardized effect sizes, and directional concordance rather than raw abundance magnitudes. Batch correction was primarily relevant to the multiplexed TMT datasets, where plex represented an identifiable technical source of variation; the label-free datasets lacked an analogous reporter-multiplex structure and were analyzed using their deposited normalization procedures in the absence of sufficiently defined batch variables.

### Elastic-net modeling and cross-validation

Elastic-net logistic regression models were trained independently for each public study to classify late versus early neuropathological stage. Late-stage disease was defined as Braak stages V–VI and early-stage disease as Braak stages 0–IV.

Predictor variables comprised the proteins from the published seven-protein plasma staging panel that were quantified within each study: NPTXR, SFRP1, APP (used as a surrogate for Aβ40), IL-13, GDNF, KLK6, and MAPT (used as a surrogate for p-tau217). APP was used as a surrogate measure for Aβ40 because it is generated through the proteolytic processing of APP, whereas APP-derived peptides can provide more readily measurable MS signals than the low-abundance Aβ40 species itself.^26^ Similarly, MAPT serves as a surrogate for p-tau217 because it is the precursor protein containing the Thr217 phosphorylation site and is often more readily quantified by bottom-up MS;^27^ moreover, MAPT and p-tau217 abundances showed strong correlations across the five studies analyzed here (**Figure S2**). Both surrogates should be interpreted in this context rather than as direct measures of the specific target species.

Because protein coverage varied across public datasets, only proteins detected within a given study were included in the corresponding model. Each protein contributed two features: its quantitative abundance and a binary indicator denoting whether the protein was detected in a given sample. Age and sex were included as covariates for all studies except PXD057298 because these metadata were unavailable. Disease status was not included as a predictor; in cohorts whose early-stage group is control-dominated, diagnosis is collinear with the late-vs-early outcome and induced complete separation.

Model performance was evaluated using nested leave-one-donor-out cross-validation to ensure complete separation of training and evaluation data. During each outer iteration, a single donor was held out for testing while all preprocessing, imputation, feature standardization, hyperparameter selection, and model fitting were performed exclusively using the remaining donors. Missing protein abundances were imputed using the minimum abundance observed within the corresponding training fold, whereas detection indicators were left unchanged.

Hyperparameters governing the elastic-net penalty were selected by stratified inner cross-validation, after which the model was refitted on the complete outer training set and used to predict the held-out donor. This process was repeated until every donor received a single out-of-sample prediction.

Classification performance was quantified using the receiver operating characteristic area under the curve (ROC-AUC) calculated exclusively from held-out predictions. To assess model robustness, we summarized the hyperparameters and model coefficients selected across outer cross-validation folds.

### Reprocessing of raw MS data with Tesorai Search

Reanalysis was performed to recover peptide- and proteoform-level information that was not available in the originally published protein quantification tables. In particular, phosphorylation was included as a variable modification to enable detection of phosphorylated tau peptides.

Raw MS files from the five studies were downloaded from the PRIDE repository and reprocessed using Tesorai Search.^28^ Search results were subsequently mapped to the corresponding donor-level sample annotations. Recovered phosphopeptides were evaluated for their detection across samples and their association with neuropathological stage. Where sufficient measurements were available, p-tau217 was incorporated into the biomarker panel and its contribution to classification performance was evaluated using the same nested cross-validation framework described above.

## Results

A recent multicenter study identified a seven-protein plasma panel, including p-tau217, that distinguished individuals with Braak V–VI from those with earlier Braak 0–IV pathology defined by tau PET.^18^ The identified panel is comprised of p-tau217, secreted frizzled-related protein 1 (SFRP1), glial cell neurotrophic factor (GDNF), interleukin 13 (IL-13), Aβ40, neuronal pentraxin receptor (NPTXR), and kallikrein-6 (KLK6). The association between this BBM panel and Braak disease stage could be influenced by multiple mechanisms, the most straightforward of which is that a change in circulating proteins in the plasma reflects perturbations within the disease brain tissue. However, mechanisms outside of the CNS, such as renal and hepatic clearance or systemic inflammation that occurs downstream of brain pathology, could also impact circulating proteins in the blood. To determine whether this recently proposed plasma staging panel reflects molecular changes occurring in the AD brain, we evaluated the panel’s performance across five independent public proteomics cohorts spanning multiple brain regions, experimental workflows, and patient populations. The datasets included 204 post-mortem human brain samples spanning AD, Parkinson’s Disease (PD), Cerebral Amyloid Angiopathy (CAA), and cognitively normal controls. Braak staging was a requirement for a study to be included in our analysis.

Other sample metadata included disease status, brain region, age, sex, post-mortem interval, and APOE genotype, though availability varied across studies. Similarly, the proteomic workflows varied and included TMT-based quantitative LC-MS/MS, DIA and label-free LC-MS/MS, and phosphopeptide enrichment. Across these cohorts, global brain proteomics identified extensive disease-associated alterations in protein abundance and phosphorylation.^20–24^

Using the blood panel, each cohort was modeled independently using nested leave-one-donor-out cross-validation to assess discrimination between early and late neuropathological stages while preventing information leakage between training and evaluation data. Despite substantial heterogeneity between studies in their cohort designs and protein measurement methods, the plasma-derived biomarker panel consistently distinguished late-stage from early-stage disease across all cohorts (**Figure 3**). Receiver operating characteristic (ROC) analysis yielded areas under the curve (AUCs) ranging from 0.78 to 1.00. Importantly, these results were obtained independently within each cohort despite differences in sample preparation, MS acquisition strategies, brain regions analyzed, and the subset of panel proteins quantified.

**Figure 3.**
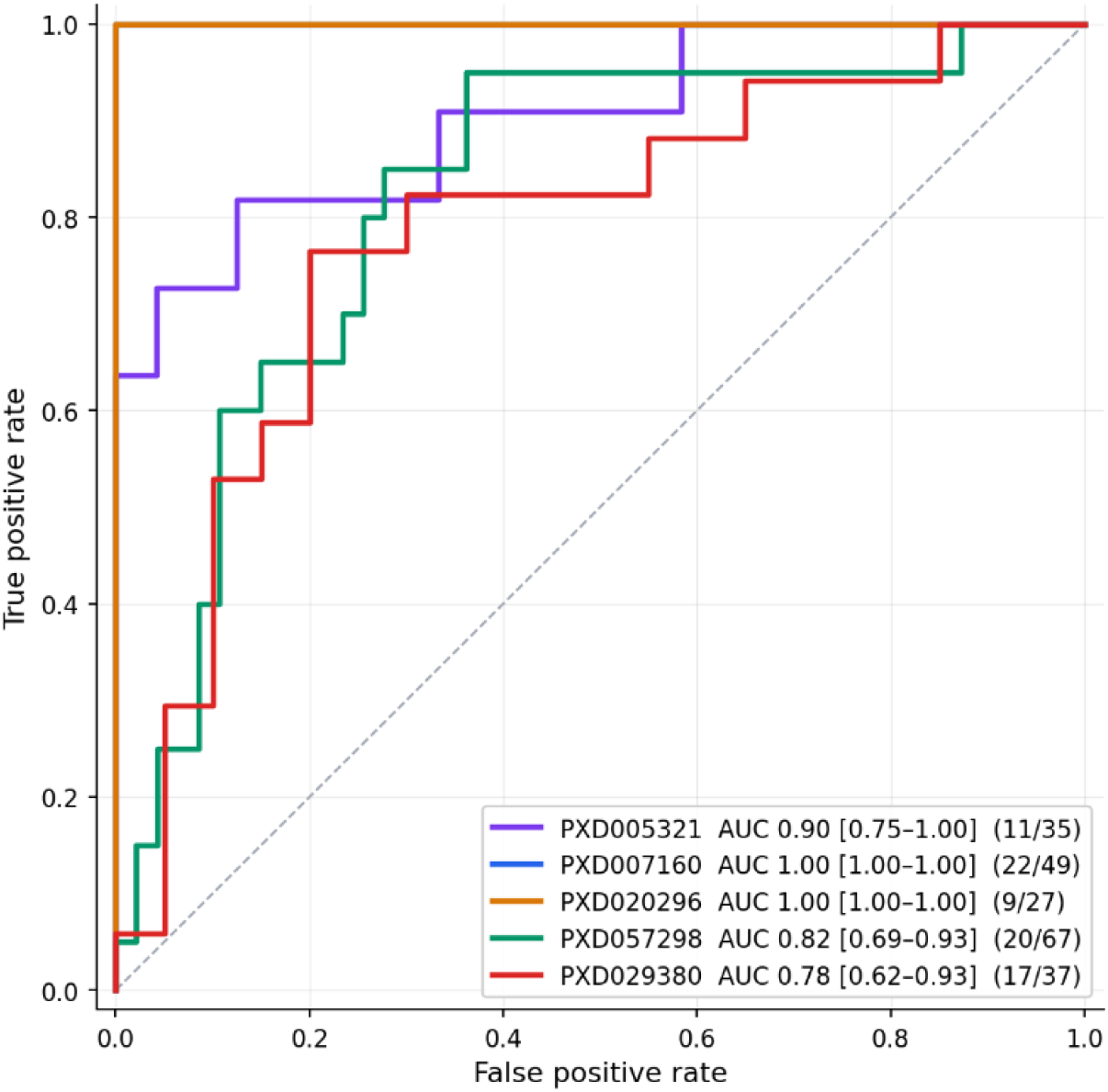
Receiver operating characteristic of the plasma tau-staging panel across five brain proteomics cohorts. Donor-level elastic-net models discriminating late-from early-stage neuropathology in each public cohort. Predictors were the measurable panel subset, NPTXR, SFRP1, APP as an Aβ40 surrogate, IL-13, GDNF, KLK6, and tau (MAPT) as a p-tau217 surrogate, with minimum-imputed abundance and a detection indicator per protein, plus age and sex where available. Each cohort was modeled independently by nested leave-one-donor-out cross-validation, with imputation and hyperparameter selection confined to training folds; curves are pooled from out-of-sample predictions. Performance is reported as area under the curve (AUC). Bracketed values are 95% confidence intervals for each AUC from 500 donor-level bootstrap resamples of the leave-one-donor-out predictions. For each study, late-stage/total donor counts are shown in parentheses. The line for PXD007160 is obscured by the line for PXD020296 due to both curves having an AUC of 1.00. Diagonal, chance (AUC 0.50).

Notably, robust classification performance was maintained even though no public dataset contained measurements for the complete plasma biomarker panel in brain tissue. Depending on the study, only two to five of the seven biomarkers were detected, reflecting the heterogeneous nature of discovery proteomics datasets. Additionally, the original analyses did not quantify disease-relevant proteoforms such as p-tau217 or the Aβ40 cleavage product. Consequently, Microtubule Associated Protein Tau (MAPT; also known as tau) and Amyloid Precursor Protein (APP) abundance were used as surrogate measurements in the present analysis.^26,27,29^ This substitution provides a reasonable proxy, as MAPT and p-tau217 abundances showed strong correlations across the five studies analyzed here (**Figure S2**). Despite this substitution, the available subset of biomarkers consistently retained substantial discriminatory power, suggesting that the molecular signature underlying the plasma staging panel is recapitulated within brain tissue rather than being a specific property of the plasma. All results were obtained independently within each cohort despite differences in sample preparation, MS acquisition, brain regions analyzed, and the subset of panel proteins detected.

The observed AUCs were higher than those achieved by randomly sampled protein panels, demonstrating discriminatory value of the panel in these cohorts (**Table S1**). The PXD005321 and PXD007160 datasets exhibited strong differentiation from random protein panels (**Table S1**). Cohort discrimination in the PXD020296 cohort reached an AUC of 1.00. However, given its small size (27 donors, 9 late-stage) and relatively high performance of random protein panels (**Table S1**), this result is best interpreted as evidence of separability rather than as a reliable estimate of generalizable classification performance. The relatively high performance of random protein panels in PXD020296 and PXD057298 likely reflects broad proteomic differences between early- and late-stage disease observed in these studies, as advanced AD is accompanied by coordinated changes across multiple biological processes, such as neuronal and synaptic function, inflammation, and cellular metabolism.^30^ Thus, assessing model performance relative to random protein panels provides an important measure of panel specificity and indicates that high AUC values should be interpreted in the context of the overall degree of proteomics separation between neuropathological stage groups (**Figure S4)**.

To assess whether individual biomarkers showed consistent associations with neuropathological stage, we fitted univariate logistic regression models for each protein and compared the resulting coefficients with those of the published plasma staging model (**Figure 4**). Four panel components, p-tau217, Aβ40, IL-13, and GDNF, were not quantified in the initial analysis of any brain cohort, restricting the comparison to MS-measurable proteins, though we were able to re-analyze raw spectra to estimate p-tau217 (see below, **Table 2**). MAPT showed the most consistent association with late-stage pathology, with positive coefficients across all informative cohorts (+0.70 to +1.91), including complete separation in PXD020296. NPTXR was negatively associated with late-stage pathology in three cohorts and positively associated in one, whereas SFRP1 and KLK6 showed weaker and more variable associations.

**Figure 4.**
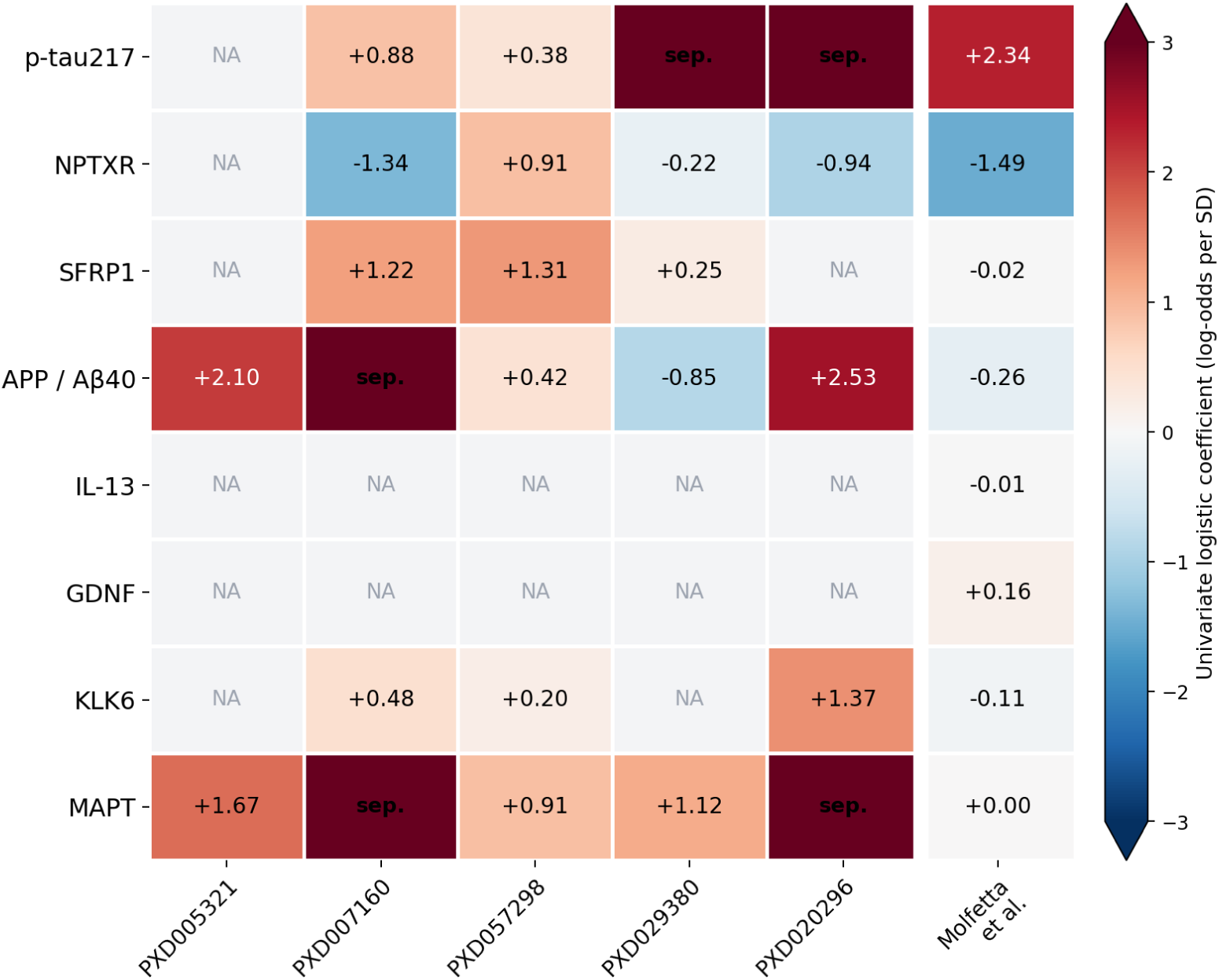
Consistency of individual biomarker associations across independent brain proteomics cohorts. Each cell represents the coefficient from a univariate logistic regression model predicting late (Braak V–VI) versus early (Braak 0–IV) neuropathological stage using a single biomarker. The rightmost column shows the corresponding coefficients from the published plasma staging model (Molfetta *et al.*). Proteins not quantified in a given study are indicated as NA. Univariate models were used to assess the direction and magnitude of each biomarker’s association independently, avoiding coefficient instability arising from collinearity in multivariable models. The consistent direction of association observed across independent cohorts supports the biological reproducibility of the published plasma biomarker panel. Sep. = complete separation.

**Table 2.**
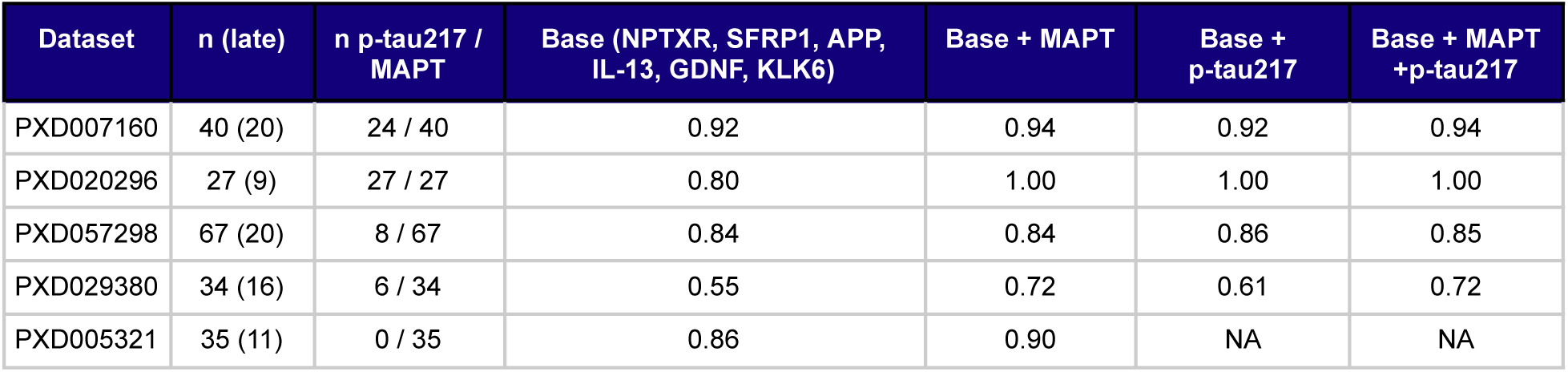
Effect of adding p-tau217 to the panel elastic-net model on discrimination of late- versus early-stage neuropathology, in the three cohorts with detectable p-tau217. The “n (late)” column is the total cohort size for that dataset’s model and the number of donors in the "late" stage neuropathology group. The “n p-tau217 / MAPT” column indicates the number of donors in which p-tau217 or MAPT was identified. The data for PXD007160 in this table restricts to the anterior-cingulate subset (n=40) and not the all-region cohort (n=49). Elastic-net models were recomputed for this table, and results for some datasets therefore differ slightly from those shown in Figure 1.

Taken together, these analyses demonstrate that the principal tau- and amyloid-related components of the published plasma staging panel exhibit reproducible associations with the AD stage across independent brain proteomics cohorts. Differences in the magnitude or direction of individual coefficients should therefore be interpreted within the context of differing cohort designs, statistical models, and measurement technologies, rather than as evidence of biological inconsistency.

### Recovery of p-tau217 from legacy proteomics datasets

Although the plasma staging panel generalized well across independent brain proteomics cohorts, none of the public datasets contained explicit measurements of p-tau217, a key component of the original panel. We therefore investigated whether disease-relevant tau phosphopeptides could be recovered directly from raw MS data despite the absence of phosphopeptide enrichment and their omission from the published protein quantification tables.

Reanalysis of the raw MS data with Tesorai Search identified the Thr217-phosphorylated tau peptide (TPSLPT[pT]PPTR) in four of the five cohorts (not detected in PXD005321). Quantifiable p-tau217 signal was observed in a subset of donors within each cohort (24/40 donors in PXD007160, 27/27 in PXD020296, and 6/34 in PXD029380) and was consistently elevated in late-stage relative to early-stage disease, demonstrating that clinically relevant phosphoproteoforms can be recovered retrospectively from unenriched discovery proteomics datasets. Notably, the AD-associated phosphopeptide p-tau231 was identified in all five cohorts and similarly showed strong associations with late Braak stage.

We next asked whether incorporating the recovered phosphopeptide into the published biomarker panel improved staging performance. To this end, we repeated the elastic-net analysis after adding p-tau217 to the feature set while leaving the model architecture and nested cross-validation procedure unchanged. Despite its central role in the published plasma panel, inclusion of p-tau217 produced only modest improvements in classification performance across the four evaluable cohorts (**Table 2**).

To determine whether the limited incremental value of p-tau217 reflected redundancy with the existing tau measurement, we performed a factorial analysis. Starting with a base panel containing only the non-tau markers (NPTXR, SFRP1, APP, and KLK6, where quantified), we added MAPT, p-tau217, or both and re-evaluated classification performance (**Table 2**). Across all cohorts, adding p-tau217 to a model already containing MAPT did not improve discrimination. In the cohort with complete p-tau217 coverage (PXD020296; p-tau217 detected in 27/27 donors), the base panel achieved an AUC of 0.80. Addition of either MAPT or p-tau217 increased performance identically to 1.00, with no further improvement when both were included, indicating that the two measurements captured largely overlapping stage-related information in this cohort. This substitution was less apparent in cohorts with incomplete p-tau217 detection. In PXD007160 (24/40 donors), MAPT provided only a modest improvement over the base panel (0.92→0.94), whereas p-tau217 did not improve performance (0.92), and including both yielded an AUC of 0.94. In PXD029380 (6/34 donors), MAPT substantially improved discrimination (0.55→0.72), whereas p-tau217 provided only a partial improvement (0.61) and added no further benefit when combined with MAPT (0.72), consistent with sparse detection limiting its contribution. Similarly, in PXD057298 (8/67 donors), the base panel already showed strong discrimination (AUC = 0.84), and neither MAPT nor p-tau217 materially changed performance (MAPT, 0.84; p-tau217, 0.86; both, 0.85). In PXD005321, where p-tau217 was not detected, MAPT nevertheless improved performance of the APP-only base panel (0.86→0.90).

Peptide-level analyses further support the interpretation that the lack of incremental benefit reflects redundancy rather than absence of disease relevance. MAPT was quantified in all three cohorts, and the stage-dependent tau signal was concentrated within peptides derived from the microtubule-binding region (MTBR), a portion of the protein that is largely devoid of disease-associated phosphorylation sites. MTBR peptides increased markedly in late-Braak donors in PXD007160 and PXD020296 (**Figure S3**). Because these peptides contribute to the aggregate MAPT abundance used by the classifier, the original panel already captured much of the tau-related signal associated with neuropathological stage, leaving limited opportunity for p-tau217 to contribute additional discriminatory power.

Taken together, these findings demonstrate that clinically relevant phosphoproteoforms can be recovered retrospectively from legacy discovery proteomics datasets and retain clear associations with AD progression. At the same time, they reveal that the incremental value of p-tau217 is tissue dependent. In brain tissue, p-tau217 staging information is largely redundant with tau abundance, whereas in plasma it provides complementary information that substantially enhances disease staging.

## Discussion

In this study, we demonstrate that a recently proposed plasma protein panel for staging advanced AD is associated with molecular remodeling in post-mortem brain tissue across multiple proteomics cohorts. We identified an opportunity to explore this hypothesis using available public datasets, but also highlight that the systematic reuse of these resources remains challenging. Brain proteomics studies differ substantially in tissue region, sample preparation, MS platform, protein inference, reporting conventions, and accompanying metadata and structure. Candidate biomarkers may therefore be present in one dataset but absent from another, even when the underlying MS data contain relevant peptide evidence. This limitation is particularly important for disease-relevant proteoforms rather than total protein abundance.

Phosphorylated tau species such as p-tau217 are commonly not represented in conventional protein-level summaries, particularly from legacy proteomics experiments. Reanalysis of raw MS data provides an opportunity to recover such disease-relevant peptides and extend the biological information available from previously generated datasets. Our results have demonstrated how pre-structuring of datasets and scalable re-analysis can rapidly establish supporting evidence for new biomarker findings.

Despite substantial heterogeneity in tissue source, experimental workflow, protein coverage, and cohort composition, the seven blood-panel proteins consistently discriminated against later neuropathological disease from earlier stages, supporting the premise that the circulating plasma signature captures biological processes directly within the AD brain. The reproducible association of MAPT with the advanced Braak stage is consistent with the established relationship between tau accumulation and disease progression. NPTXR is particularly notable given its emerging role as a marker of synaptic integrity and neurodegeneration.^31^ Recent longitudinal studies have shown that reduced CSF NPTXR tracks cognitive impairment, cortical thinning, and subsequent clinical progression, further supporting its interpretation as a marker of downstream neurodegenerative biology rather than amyloid pathology alone.^32^ Other components of the panel, including KLK6 and SFRP1, may capture complementary biological processes involving neurodegeneration, proteolysis, and amyloid-associated signaling, although the current evidence for their contributions to disease staging is less established and may be dependent on biological compartment and disease context. In particular, the heterogeneous directionality reported for KLK6 across brain, CSF, and plasma highlights the importance of distinguishing molecular measurements in the tissue of disease from their circulating counterparts.^33^

An additional finding was that reanalysis of legacy discovery proteomics data with Tesorai Search recovered the disease-relevant p-tau217 phosphoproteoform in multiple cohorts despite its absence from conventional protein-level summaries. Although p-tau217 was strongly associated with advanced neuropathological stage, its addition to brain-based models provided little incremental discrimination when MAPT was already included, consistent with substantial overlap between phospho-tau and tissue tau abundance in the brain. Thus, these findings underscore that the biological information contributed by a biomarker is inherently dependent on both the molecular species measured and the biological compartment in which it is measured.

One limitation of our approach is that protein coverage differs across datasets. For example, two proteins, IL-13 and GDNF, were not detected in any of the brain datasets. The absence of these proteins may reflect their low abundance and sensitivity limits of the technology rather than a true biological absence. Supporting this, both of these proteins have been detected in brain tissue of AD patients using targeted, more sensitive approaches.^34–39^

Several coefficients, particularly APP/Aβ40 and NPTXR, differed in direction from the published plasma model. Although Aβ42 is well established to decline in the plasma with increasing amyloid plaque burden in the brain, Aβ40 has been reported to remain relatively stable.^40,41^ Both Aβ42 and Aβ40 are non-trivial to measure reliably, as discussed by di Molfetta *et al*.^18^ As such, it is likely that the observed differences in coefficient direction reflect a combination of statistical, analytical, and biological factors: the plasma model estimates multivariable associations, whereas our analysis evaluates marginal effects; immunoassays and MS quantify different molecular species; and plasma and brain represent distinct biological compartments.

Additionally, the relatively strong performance of randomly selected protein panels in some cohorts also limits interpretation of AUC as a generalizable measure. Broad proteomic remodeling associated with advanced neuropathological disease can result in many proteins carrying information about disease stage, particularly in small cohorts. Thus, the present analyses support reproducibility of stage-associated information within the proposed panel but do not establish that the panel is uniquely predictive relative to the broader brain proteome.

Recovery of p-tau217 from legacy datasets was also limited by the characteristics of the original acquisition workflows. The phosphopeptide was detected in only a subset of cohorts and donors, and the absence of phosphopeptide enrichment and differences in acquisition depth likely reduced sensitivity. Consequently, nondetection cannot be interpreted as biological absence, and the incremental performance of p-tau217 could not be evaluated consistently across all cohorts.

Finally, additional limitations are that the analyzed cohorts differed in the diagnosed disease or cohort definition, anatomical regions sampled, and post-mortem interval, all of which may influence both baseline protein abundance and the magnitude of disease-associated changes. Because AD pathology progresses in a regionally heterogeneous manner and tissue quality can impact protein measurement, these factors may contribute to between-cohort variation in biomarker associations and model performance.

Our results support recently identified multiprotein blood-based signatures^18^ as indicators of advanced AD pathology. More broadly, they demonstrate the value of systematically re-mining legacy proteomics datasets to establish tissue-level biological concordance, test biomarker generalizability across independent cohorts, and recover disease-relevant proteoforms that may be absent from conventional protein-level summaries. Prospective studies integrating standardized plasma and CSF measurements with amyloid and tau imaging, neuropathology, cognition, and longitudinal outcomes will be important to determine how individual components of the panel contribute to staging, prognosis, and treatment-response monitoring.

## Supplemental Material

**Figure S1:**
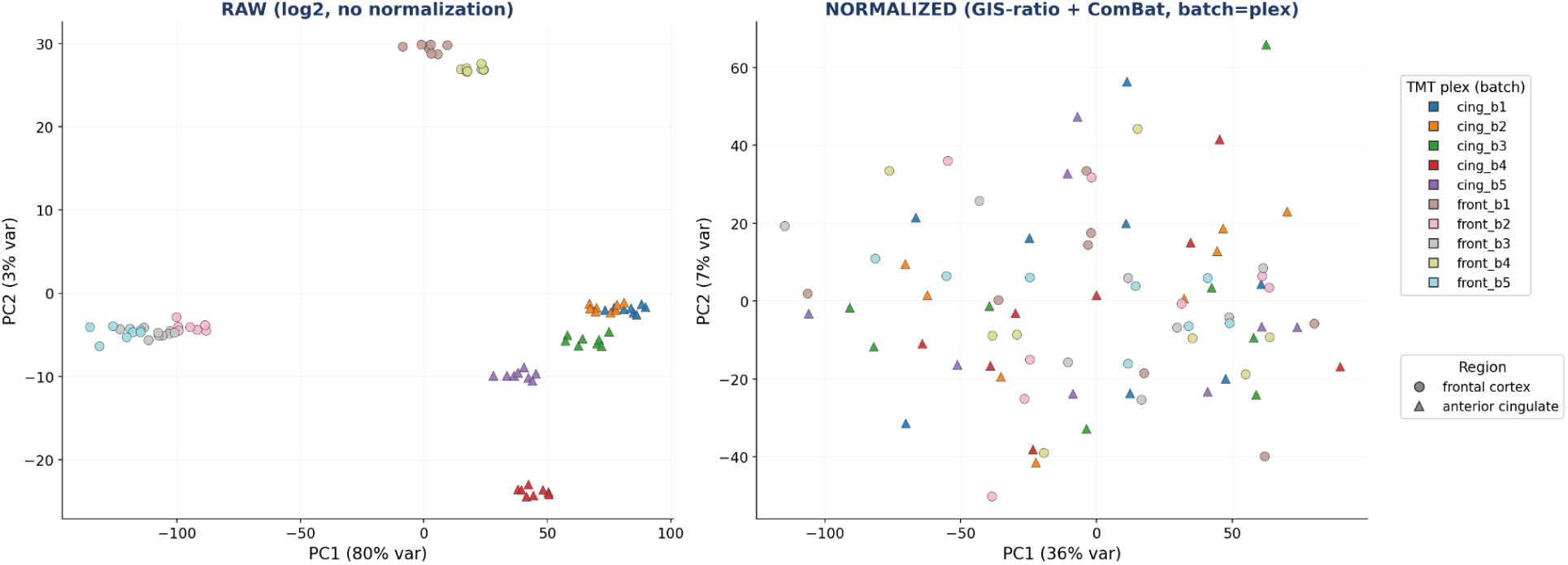
Removal of TMT batch effects in PXD007160 by GIS-ratio normalization and ComBat. PCA analysis of individual specimens before (left) and after (right) normalization, with each point a specimen colored by TMT plex (batch) and shaped by brain region. In the raw data, specimens cluster tightly by plex and the first principal component is dominated by batch; after within-plex normalization to the pooled reference channels followed by ComBat batch correction, the plex clusters dissolve and batch-associated variance is largely removed (PC1–plex association R² 12.9% → 0.04%). Percentages on each axis indicate the proportion of variance explained.

**Figure S2.**
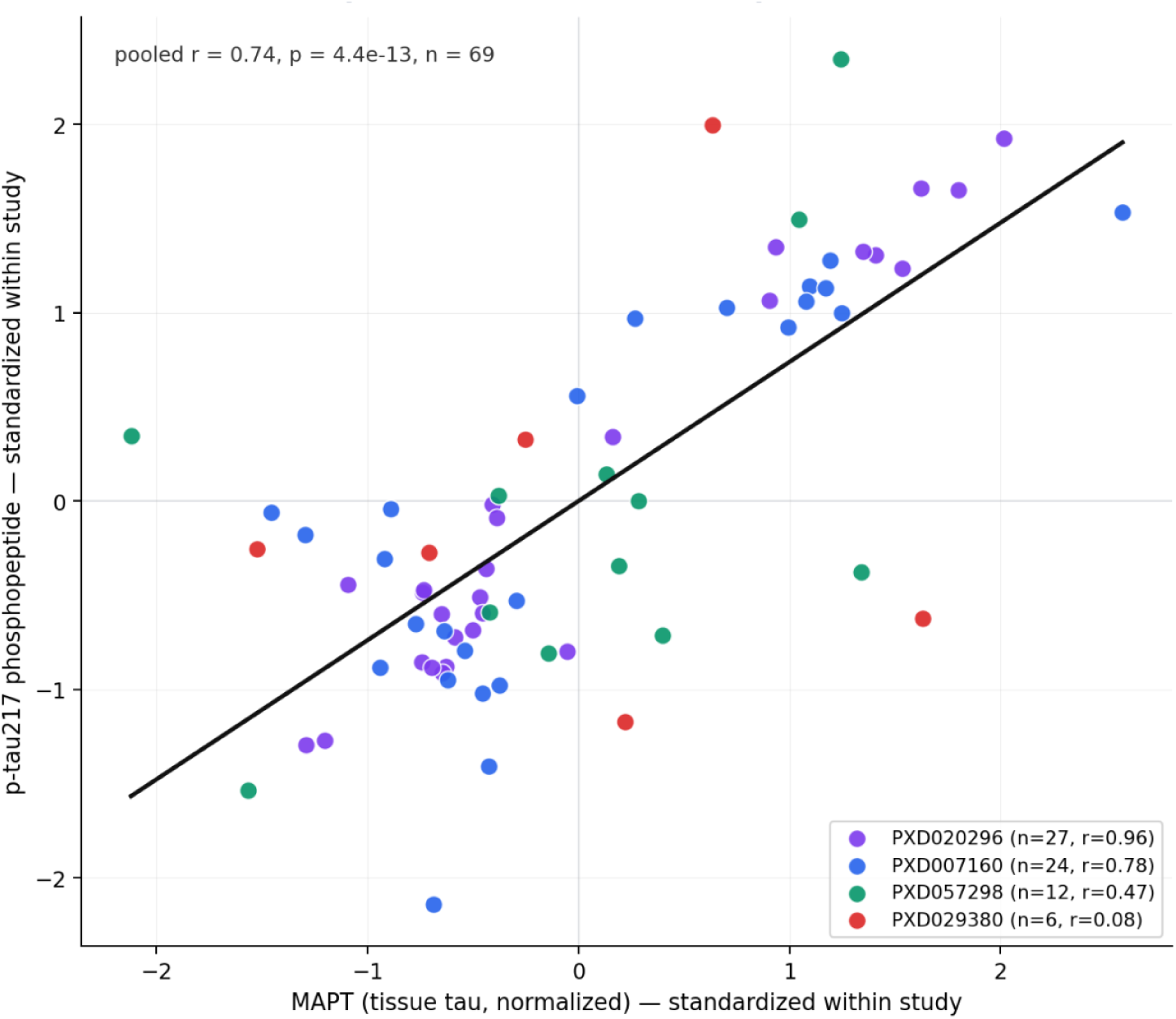
p-tau217 covaries with total tissue MAPT across cohorts. Per-donor levels of p-tau217 plotted against normalized MAPT abundance in the four datasets where p-tau217 was detected. Both axes are standardized (z-scored) within each study to allow pooling across platforms (TMT reporter ratios, label-free precursor, DIA); each point is one donor, colored by study. The black line represents the pooled linear fit. p-tau217 and MAPT are positively correlated (pooled r = 0.74, p ≈ 4×10⁻¹³, n = 69; within-study r = 0.96, 0.78, 0.47, and 0.08 for PXD020296, PXD007160, PXD057298, and PXD029380 respectively).

**Figure S3.**
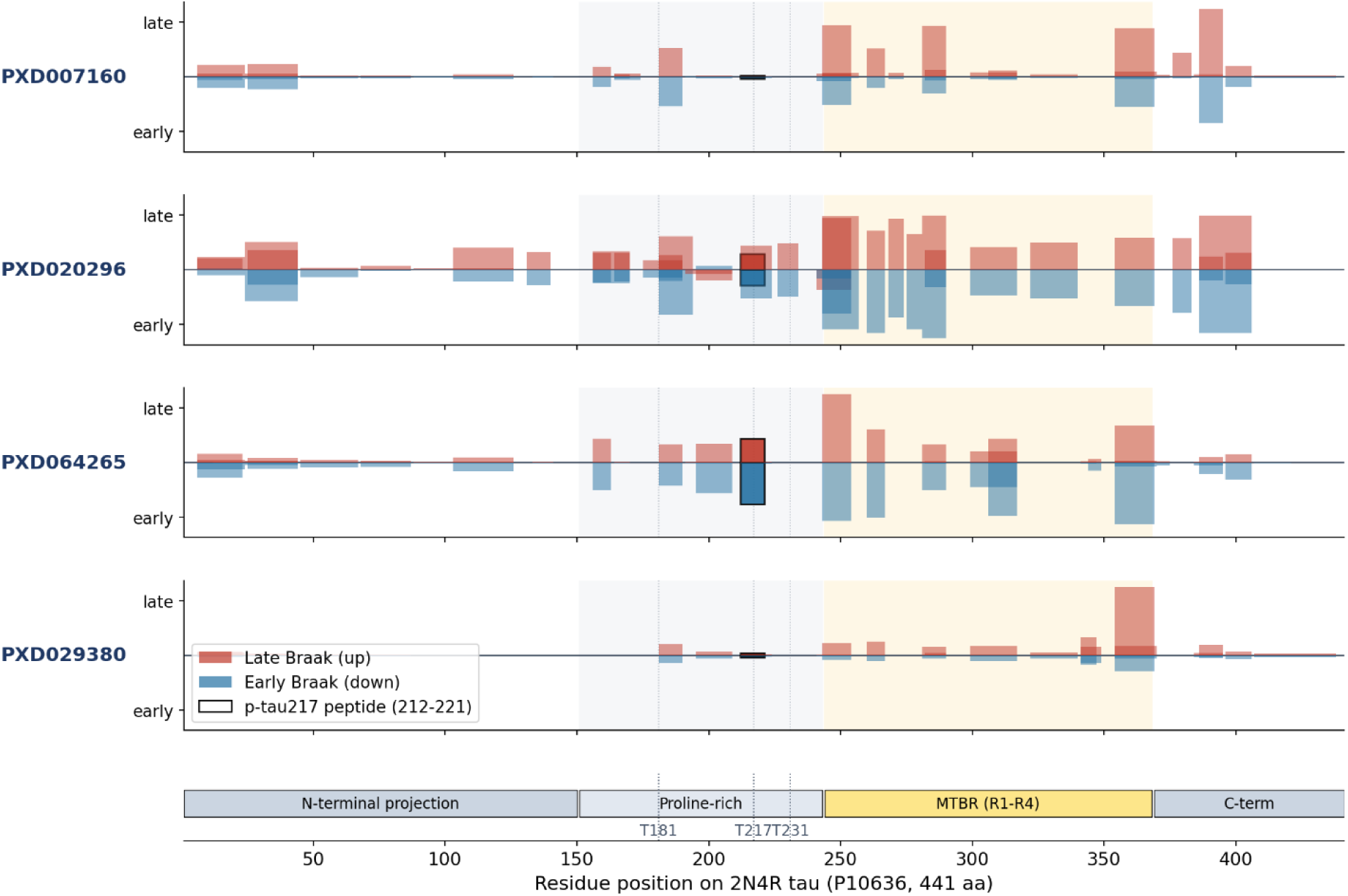
MAPT peptide intensities across the tau protein stratified by Braak stage. Upward and downward bars represent peptides that are more abundant in late and early Braak stage samples, respectively. Bar thickness corresponds to the span of the peptide along the MAPT protein sequence. Peptide abundances are reproduced from the original published datasets and were not generated by our reanalysis. Consistent with this interpretation, the corresponding unmodified peptide spanning residues 212–221 (TPSLPTPPTR) showed little or no stage-dependent difference across cohorts, indicating that the discriminative signal arises from phosphorylation at Thr217 rather than changes in peptide abundance alone. Thus, while p-tau217 provides complementary information in plasma, where tau is only modestly informative, it contributes comparatively little in brain tissue, where overall tau accumulation is already strongly associated with disease stage. In PXD029380, where p-tau217 was detected in only a small fraction of donors, its inclusion marginally reduced classification performance, consistent with sparse measurements contributing more noise than signal.

**Figure S4.**
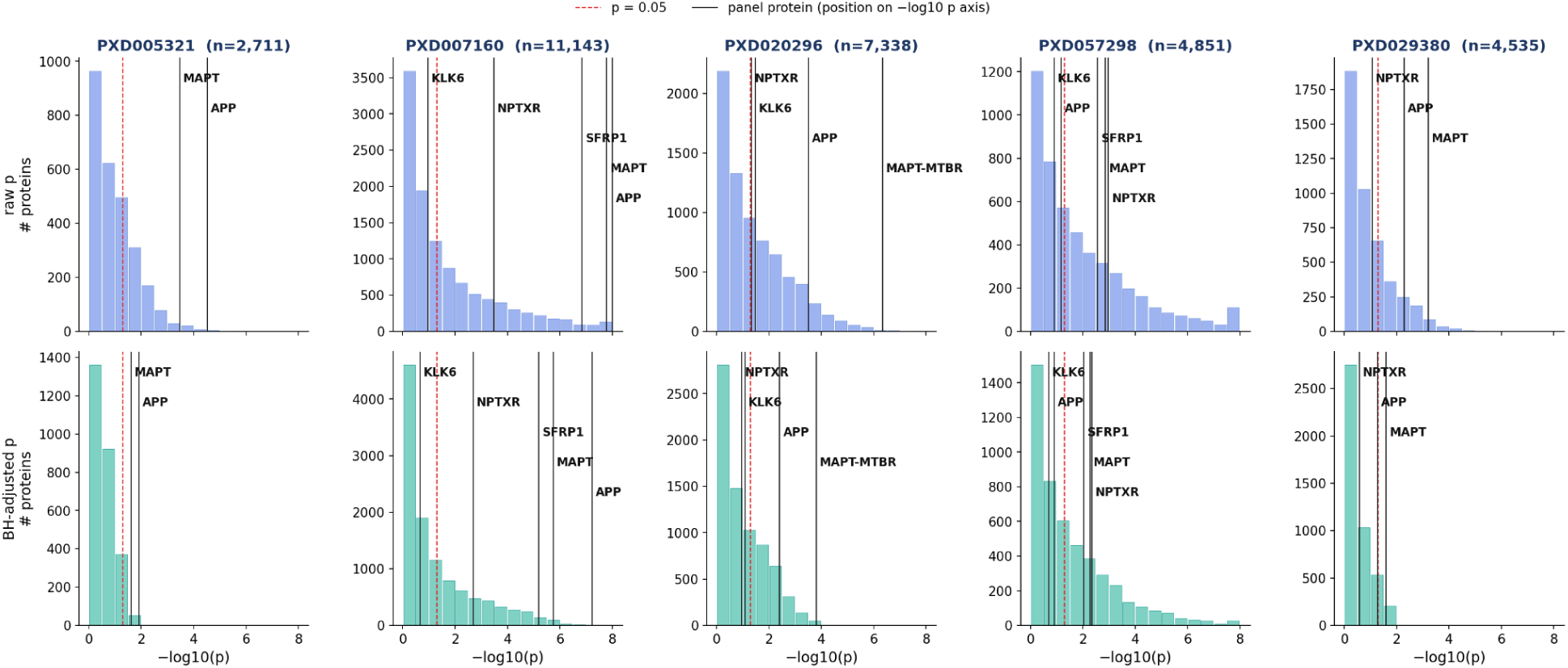
Significance of individual panel proteins. Histograms of p-values and Benjamini-Hochberg adjusted p-values are shown for each study, with the significance of individual panel proteins labeled.

**Table S1.** Random-protein negative control for panel-based Braak staging. For each cohort, six proteins were drawn uniformly at random from the full quantified proteome and used to predict late- versus early-stage neuropathology (Braak V–VI vs 0–IV, using the same donor-level pipeline as the biomarker panel (minimum-imputed abundance plus a detection indicator per protein, age and sex as covariates where available, elastic-net logistic regression evaluated by pooled leave-one-donor-out cross-validation). The draw was repeated 10 times per cohort (seeds 1–10); the mean, minimum, and maximum out-of-sample AUC across draws are shown alongside the biomarker-panel AUC for comparison. In PXD005321 and PXD007160 all random draws performed below the panel, indicating cohort-specific predictive value; in PXD020296 and PXD057298 random draws frequently matched or approached the panel performance, indicating broad separability between stage groups (consistent with a diagnosis-driven global proteome difference) rather than panel-specific signal. Proteins were sampled uniformly over all quantified protein groups, so individual draws could include tau- or panel-adjacent proteins. AUC, area under the receiver operating characteristic curve.

| Dataset | k | Random-k mean (range) | Panel AUC |
| --- | --- | --- | --- |
| PXD005321 | 2 | 0.67 (0.59–0.77) | 0.90 |
| PXD007160 | 5 | 0.67 (0.23–0.94) | 1.00 |
| PXD029380 | 4 | 0.67 (0.36–0.81) | 0.78 |
| PXD020296 | 4 | 0.75 (0.40–0.96) | 1.00 |
| PXD057298 | 5 | 0.76 (0.61–0.85) | 0.82 |

